# Probe capture enrichment sequencing of *amoA* genes discloses diverse ammonia-oxidizing archaeal and bacterial populations

**DOI:** 10.1101/2023.04.10.536224

**Authors:** Satoshi Hiraoka, Minoru Ijichi, Hirohiko Takeshima, Yohei Kumagai, Ching-Chia Yang, Yoko Makabe-Kobayashi, Hideki Fukuda, Susumu Yoshizawa, Wataru Iwasaki, Kazuhiro Kogure, Takuhei Shiozaki

**Affiliations:** Research Center for Bioscience and Nanoscience (CeBN), Japan Agency for Marine-Earth Science and Technology (JAMSTEC), 2–15 Natsushima-cho, Yokosuka, Kanagawa 237–0061, Japan; Atmosphere and Ocean Research Institute, the University of Tokyo, 5-1-5 Kashiwanoha, Kashiwa, Chiba 277-8564, Japan; Department of Integrated Biosciences, Graduate School of Frontier Sciences, the University of Tokyo, 5-1-5 Kashiwanoha, Kashiwa, Chiba 277-0882, Japan

**Keywords:** Ammonia-oxidizing prokaryotes, *amoA* gene, Target gene enrichment, Hybridization capture, Metagenome, Metatranscriptome, Marine prokaryotes

## Abstract

The ammonia monooxygenase subunit A (*amoA*) gene has been used to investigate the phylogenetic diversity, spatial distribution, and activity of ammonia-oxidizing archaeal (AOA) and bacterial (AOB), which contribute significantly to the nitrogen cycle in various ecosystems. Amplicon sequencing of *amoA* is a widely used method; however, it produces inaccurate results owing to the lack of a ‘universal’ primer set. Moreover, currently available primer sets suffer from amplification biases, which can lead to severe misinterpretation. Although shotgun metagenomic and metatranscriptomic analyses are alternative approaches without amplification bias, the low concentration of target genes in heterogeneous environmental DNA restricts a comprehensive analysis to a realizable sequencing depth. In this study, we developed a method for *amoA* enrichment sequencing using a hybridization capture technique. Using metagenomic mock community samples, our approach effectively enriched *amoA* genes with low compositional changes, outperforming amplification and meta-omics sequencing analyses. Following the analysis of metatranscriptomic marine samples, we predicted 80 operational taxonomic units (OTUs) assigned to either AOA or AOB, of which 30 OTUs were unidentified using simple metatranscriptomic or *amoA* gene amplicon sequencing. Mapped read ratios to all the detected OTUs were significantly higher for the capture samples (50.4 ± 27.2%) than for non-capture samples (0.05 ± 0.02%), demonstrating the high enrichment efficiency of the method. The analysis also revealed the spatial diversity of AOA ecotypes with high sensitivity and phylogenetic resolution, which are difficult to examine using conventional approaches.

## Introduction

Nitrification is a two-step process involving ammonia oxidation followed by nitrite oxidation. Ammonia oxidation is a rate-limiting process driven by ammonia-oxidizing archaea (AOA) and bacteria (AOB), which utilize the ammonia monooxygenase subunit A (*amoA*) gene. This process is fundamental to the nitrogen cycle, including the emission of nitrous oxide, a primary greenhouse gas (Fowler et al., 2013; Niu et al., 2016), and occurs in various ecosystems such as soil, hot springs, freshwater, estuaries, sediments, and marine ecosystems (Isobe & Ohte, 2014; Lehtovirta-Morley, 2018; Monteiro et al., 2014; Takai, 2019). A substantial proportion of the ammonia oxidation process on Earth is believed to occur in the ocean via AOAs (Hutchins & Capone, 2022; Voss et al., 2013). Oceanic AOAs are largely classified into three ecotypes: water column cluster A (WCA), water column cluster B (WCB), and *Nitrosopumilus maritimus*–like cluster (NMC, known as NPM) (Beman et al., 2008; Francis et al., 2005; Santoro et al., 2019). WCA and NMC have been reported to be abundant just below the euphotic zone, whereas WCB inhabits deeper waters (Beman et al., 2008; Santoro et al., 2019; Shiozaki et al., 2016). According to the recently updated taxonomy based on extensive AOA *amoA* gene phylogeny (Alves et al., 2018), WCA, WCB, and NMC correspond to NP-ε-2, NP-α-2.2.2.1, and NP-γ-2.1, respectively. Although AOAs contribute greatly to ammonia oxidization in general, AOBs are compatible with AOAs in environments with high substrate availability (e.g., agricultural soil), regardless of their lower relative abundance (Jia & Conrad, 2009; Prosser et al., 2020; Sterngren et al., 2015). Among the AOBs, complete ammonia oxidizers (comammox) can perform the entire two-step nitrification process alone, are found in both terrestrial and coastal areas, and are considered important contributors to nitrification on a global scale (Daims et al., 2015; Fei et al., 2018; van Kessel et al., 2015).

*amoA* amplicon sequencing is a commonly used approach for characterizing AOAs and AOBs in prokaryotic communities. This methodology relies on polymerase chain reaction (PCR) for the selective amplification of *amoA* without cultivation. Currently, *amoA* is the second most frequently sequenced marker gene in microbiology, preceded by the 16S ribosomal RNA (rRNA) gene (Alves et al., 2018), allowing us to investigate the phylogenetic diversity and spatial distribution of diverse AOA and AOB in various environments, including the ocean (Table S1) (Beman et al., 2008; Francis et al., 2005; O’Mullan & Ward, 2005). However, PCR is not an all-encompassing technique. First, it is difficult to design a universal primer set for all AOA and AOB owing to the lack of a conserved region in the cording sequence of *amoA* gene. Although a number of primer sets with different targeted clades have been proposed (Beman et al., 2008; Coolen et al., 2007; Francis et al., 2005; Hallam et al., 2006; Hornek et al., 2006; Leininger et al., 2006; Löscher et al., 2012; Mincer et al., 2007; Mosier & Francis, 2011; Pester et al., 2012; Pjevac et al., 2017a, 2017b; Purkhold et al., 2000; Rotthauwe et al., 1997; Sintes et al., 2013; Wuchter et al., 2006; Yakimov et al., 2011), a true ‘universal’ primer set covering both AOA and AOB is yet to be developed. Secondly, even among the targeted clades of the primer set used, PCR amplification frequently causes unexpected results owing to mismatches between the primer and targeted gene sequences, differences in the efficiency of the denaturation and annealing steps, and chimeric amplification, known as PCR biases (Jolinda et al., 2018). For example, the most frequently used *amoA* primer set, Arch-amoAF/R (Francis et al., 2005), results in mismatches with numerous AOA *amoA* sequences, which further leads to the underestimation of total AOAs (Ijichi et al., 2019; Tolar et al., 2013) and is unable to amplify any AOB *amoA* gene. Taken together, the current *amoA* amplicon sequencing approach produces distorted results, leading to misinterpretation of downstream analyses (Dechesne et al., 2016; Ijichi & Hamasaki, 2017; Shimomura et al., 2012).

Shotgun metagenomics and metatranscriptomics employ high-throughput sequencing technology to generate community-wide sequencing reads without PCR amplification. These strategies allowed us to analyze diverse gene sequences to estimate the original lineages and expression levels of each gene of interest, thereby providing a complete picture of taxonomic diversity and activity (Hiraoka et al., 2016). However, the target genes typically exist at low concentrations in heterogeneous DNA samples. The concentration of *amoA* was estimated to be up to 10^8^ copies per ng of DNA in soil (Bahram et al., 2022; Bannert et al., 2011; Ding et al., 2020; Han et al., 2013; Marusenko et al., 2013), wastewater (Spasov et al., 2020; Wang et al., 2021; Zhang et al., 2015), estuaries (Hollibaugh et al., 2011), and seawater (Christman et al., 2011) using quantitative real-time PCR (qPCR), and less than 0.001% and 0.05% of the metagenomic and metatranscriptomic reads, respectively, were assigned to the *amoA* gene from soil samples (Bahram et al., 2022; Clark et al., 2021; Paungfoo-Lonhienne et al., 2017). This low concentration limits the community-wide analysis of genes of interest to a realizable sequencing depth.

Targeted gene enrichment is an innovative approach that overcomes the limitations of PCR amplification and meta-omics. This method relies on solution hybrid selection (SHS) to recover genes of interest from heterogeneous DNAs for sequencing. Hybridization authorizes mismatches between the designed probes and distant targeted sequences, allowing for the investigation of large pools of divergent genes, including previously unknown ones (Hedtke et al., 2013). The hybridization capture experiments demonstrated high reproducibility (Chilamakuri et al., 2014; Gnirke et al., 2009; Hernandez-Rodriguez et al., 2018; Mamanova et al., 2010). In field research, this method has been applied to various genes as biomarkers, including methyl coenzyme M reductase subunit A (Denonfoux et al., 2013), endo-xylanase (Bragalini et al., 2014), carbohydrate-active enzyme and protease (Manoharan et al., 2015), reductive dehalogenase, insertion elements, and hypothetical proteins from *Dehalococcoidia* (Biderre-Petit et al., 2016), benzylsuccinate synthase alpha subunit (Ranchou-Peyruse et al., 2017), lindane degradation enzyme (Gasc & Peyret, 2018), CH_3_Cl dehalogenase (Jaeger et al., 2018), hexachlorocyclohexane degradation enzyme (Gasc & Peyret, 2017), disease resistance genes from plants (Jupe et al., 2013), 16S rRNA (Beaudry et al., 2021; Biderre-Petit et al., 2016; Cariou et al., 2018; Gasc & Peyret, 2018; Günther et al., 2022; Sara et al., 2020), and 18S rRNA(Günther et al., 2022). A pioneering study also attempted the enrichment sequencing of *amoA* gene recently (Siljanen et al., 2022). However, to the best of our knowledge, no study has focused on *amoA* probe capture enrichment sequencing with near-full-length sequence reconstruction and *de novo* identification of operational taxonomic units (OTUs) for precise analysis of phylogenetic diversity, despite their great importance in the nitrogen cycle.

Here, we developed a probe capture enrichment sequencing method for effective analysis of *amoA* genes in metagenomic and metatranscriptomic samples. Capture probes were designed using *amoA* gene sequences from a wide range of prokaryotes retrieved from public databases to cover diverse *amoA* genes, including novel ones that would hybridize with the probes. Technical validation using an artificial mock community showed the high efficiency of the approach without severe compositional disorder. The analysis of metatranscriptomic marine samples collected in the euphotic to mesopelagic layers from coastal to open ocean areas (Figure S1, Table S2) showed outstanding performance compared to conventional shotgun and amplicon sequencing approaches. The analysis also showed that the spatial distribution of AOA ecotypes could be explained by niche separation through the vertical water column and distance from the land, providing insights into nitrogen cycling in the oceanic environment and highlighting the advantages of the *amoA* enrichment sequencing approach.

## Materials and methods

### Capture probe design

The *amoA* gene belongs to the copper membrane monooxygenase (CuMMO) superfamily, which comprises several sequentially close but functionally different genes (e.g., *amoA*, *pmoA*, *pxmA*, *bmoA*, *peoA*, *etnA*, *pamA*, *AlkB*, *and bmoX*) that accept different substrates (e.g., ammonia, methane, propene, ethene, and butane). CuMMOs exhibit complicated phylogenies; thus, it is challenging to distinguish them based on sequence similarity (Alves et al., 2018; Coleman et al., 2012; Diamond et al., 2022; Orellana et al., 2016; Tavormina et al., 2011). To design capture probes that cover a wide range of *amoA* genes, we designed probes to capture diverse CuMMO genes, allowing manual selection of potential *amoA* genes for downstream sequence analysis. We manually collected 20 well-annotated representative CuMMO subunit A genes spanning a wide range of phylogenetic clades retrieved from the NCBI database as the initial database (Data S1). The sequences include four *amoA* genes from all the major AOA and AOB groups (Thaumarchaeota, Betaproteobacteria, Gammaproteobacteria, and Nitrospirota). Using the initial database, 29683 sequences were retrieved from NCBI nt and env_nt (March 2016) using blastn search with a cut-off e-value ≤ 1E-5, and 195 *amoA* gene sequences were manually collected from GenBank (Data S2). In total, 29878 CuMMO gene sequences (referred to as the CuMMO gene sequence database) were used to design 100-mer probes using a tiling strategy. To prevent the probe design from capturing an excessive amount of sequences shared by multiple genes, all duplicated sequences were eliminated with the exception of one copy. In addition, to increase the recruitment power of metatranscriptome samples, sequences that showed similarity to the 16S rRNA gene in the Ribosomal Database Project (RDP) database (Cole et al., 2014) were removed. The final probe set represented 97.06% of the total nucleotides in the CuMMO sequences. Probe design and production were performed by Roche NimbleGen (Madison, WI, USA), which is currently transferred to Roche Diagnostics (Rotkreuz, Switzerland).

### Mock sample construction and DNA sample preparation

To examine the conditions and efficiency of our probe-capture enrichment method, we prepared an artificial mixture of six cloning vectors encoding different *amoA* genes. Six *amoA* sequences were selected from three AOA and three AOB lineages: *Nitrosopumilus maritimus* SCM1^T^ (CP000866), *Candidatus* (*Ca.*) Nitrosopelagicus brevis CN25 (CP007026), Thaumarchaeota archaeon SCGC AAA007-O23 (ARWO00000000), *Nitrosomonas europaea* ATCC 19718^T^ (AL954747), *Nitrosococcus oceani* ATCC 19707^T^ (NC_007484), and *Ca.* Nitrospira inopinata ENR4 (LN885086) Six sequences were artificially synthesized by Eurofins Genomics (Ota, Tokyo, Japan) and cloned into the pTAKN-2 vector (BioDynamics Laboratory, Bunkyo, Tokyo, Japan). Each cloning vector was transformed into *Escherichia coli* DH5α competent cells (Takara Bio, Kusatsu, Shiga, Japan) according to the manufacturer’s instructions. *E. coli* was cultured in LB liquid medium supplemented with kanamycin, and cloning vectors were extracted using the PureLink HiPure Plasmid FP (Filter and Precipitator) Maxiprep Kit (Invitrogen, Carlsbad, CA, USA) following the manufacturer’s instructions. The quality and quantity of the plasmid DNA were measured using a NanoDrop 2000 (Thermo Fisher Scientific, Wilmington, DE, USA) and a Microplate Reader SH-9000Lab (Corona Electric, Hitachinaka, Ibaraki, Japan) with a Quant-iT PicoGreen dsDNA Assay Kit (Invitrogen) following the manufacturer’s instructions.

### Seawater sampling and RNA sample preparation

Seawater samples were collected from five and six stations along the Otsuchi (OT) and Onagawa (ON) transect lines (from the coastal area inside each bay to the pelagic area), respectively, during cruise KS16-01 of the R/V Shinsei-maru in March 2016 (Table S1, Figure S1). The sampling stations are located in the temperate western North Pacific Ocean. Sampling permits for expeditions in Japan’s exclusive economic zones were not required because our study was centered on domestic areas and did not include endangered or protected species. Seawater was collected using a bucket from the surface (0 m below sea level [mbsl]) or using Niskin bottle samplers (General Oceanic, Miami, Florida, USA) in deeper layers, ranging from 33 to 500 mbsl (epipelagic to mesopelagic zones). Water temperature and salinity in the sampling layer were measured using an SBE9plus CTD system (Sea-Bird Scientific, Bellevue, WA, USA). For the molecular experiments, approximately 8 L of seawater was filtered onto 0.22-μm Sterivex-GP pressure filter units (Millipore, Billerica, MA, USA). The filters were immediately soaked in 2 mL of RNA*later* Stabilization Solution (Thermo Fisher Scientific) and maintained at 4 °C for 12 h of immersion. The filters were then stored at −30 °C until subsequent molecular experiments. We also collected samples for nutrient analyses in duplicate in 10-mL acrylic tubes.

RNA was extracted from the filters onshore using the *mir*Vana miRNA Isolation Kit (Thermo Fisher Scientific), purified using the TURBO DNA-*free* Kit (Thermo Fisher Scientific), and concentrated using the RNeasy MinElute Cleanup Kit (QIAGEN, Hilden, Germany). The quality and quantity of RNA were measured using a Nanodrop 2000 and Quantus fluorometer with the QiantiFluor RNA System (Promega, Madison, WI, USA), following the manufacturer’s instructions. Concentrations of nitrate (NO_3_ ^−^), nitrite (NO_2_ ^−^), ammonium (NH_4_ ^+^), and phosphate (PO_4_ ^3−^) were measured colorimetrically using an autoanalyzer as described previously (Fukuda et al., 2016). The detection limits of the used system were 0.05, 0.02, 0.1, and 0.02 µM for NO_3_ ^−^, NO_2_ ^−^, NH_4_ ^+^, and PO_4_ ^3−^ measurements, respectively.

### Library construction, probe capture, and sequencing

Sequencing library preparation and probe capture enrichment were performed according to the SeqCap EZ Developer Kit (Roche NimbleGen) or the SeqCap RNA Developer Enrichment Kit (Roche NimbleGen). Before library construction, the quality of all RNA samples was verified using a 2100 Bioanalyzer (Agilent Technologies) and an RNA Pico Kit (Agilent Technologies), which confirmed that no severe RNA degradation was observed. DNA and RNA were shared to a size of 200 bp via sonication using the Covaris (Woburn, MA, USA) and exposure to divalent cations in accordance with the SeqCap kits. cDNA was synthesized from RNA using a KAPA Stranded RNA-Seq Library Preparation Kit (Roche NimbleGen). DNA/cDNA libraries with index adapters were then prepared using the KAPA Library Preparation Kit (Roche NimbleGen) and the KAPA Stranded RNA-Seq Library Preparation Kit (Roche NimbleGen) according to the manufacturer’s instructions.

For probe capture enrichment sequencing, hybridization capture was performed using the SHS method according to the SeqCap EZ Reagent Kit Plus (Roche NimbleGen) with modifications to the number of hybridization captures and the number of post-capture ligation-mediated PCR (LM-PCR) cycles (see below). The basic protocol for probe capture enrichment was as follows: The DNA/cDNA libraries were LM-PCR amplified using KAPA HiFi HotStart ReadyMix according to the official instructions (pre-capture LM-PCR): 98 °C for 45 s, (98 ℃ for 15 s, 60 ℃ for 30 s, 72 ℃ for 30 s) cycle for 9 times, 72 ℃ 1 min for DNA libraries, and 98 ℃ for 45 s, (98 ℃ for 15 s, 60 ℃ for 30 s, 72 ℃ for 30 s) cycle for 11 times, 72 ℃ for 5 min for cDNA libraries. After purification, equimolar libraries were mixed to prepare a library pool for each analysis (details are provided below). Next, 1 μg of the library pool was denaturated at 95 ℃ for 10 min and subjected to probe hybridization at 47 ℃ for 16–20 h. The probes were captured by SeqCap Capture Beads at 47 ℃ for 45 min with Vortex shaking every 15 min. Then, the bead-bound capture DNA/cDNA libraries after washing were directly LM-PCR amplified using KAPA HiFi HotStart ReadyMix (post-capture LM-PCR): 98°C for 45 s, (98 ℃ for 15 s, 60 ℃ for 30 s, 72 ℃ for 30 s) cycle for 20 times, 72 ℃ for 1 min for DNA libraries, and 98°C for 45 s, (98 ℃ for 15 s, 60 ℃ for 30 s, 72 ℃ for 30 s) cycle for 20 times, 72 ℃ for 5 min for cDNA libraries. After purification, amplified libraries were sequenced. The pre- and post-capture LM-PCR amplifications were originally incorporated into the official instructions of the SeqCap kits to obtain sufficient amounts of DNA for hybridization capture and sequencing, respectively.

To evaluate potential biases in probe capture enrichment and to optimize the methodology, different settings were tested using the mock sample and a seawater sample collected from 0 mbsl at St2 station (Table 1). In this experiment, the times of probe capture enrichment were varied with zero, once, and twice, referred to as ‘non-capture,’ ‘single-capture,’ and ‘double-capture’ settings, respectively. For the non-capture setting, libraries were prepared without probe capture enrichment, thus indicating an unenriched shotgun sequencing that can produce a bias-free community composition. For single-capture settings, probe capture enrichment was performed once. For the double-capture settings, probe capture enrichment was performed twice with different combinations of the first and second post-capture LM-PCR amplifications (pairs of 5 and 20 (5/20), 7 and 14 (7/14), and 7 and 20 (7/20) cycle settings for the mock sample and 5/20 and 7/14 cycle settings for the seawater sample). For the probe capture enrichment, the libraries from Mock and St2 samples were mixed as a library pool, 1 μg of which was subjected to probe hybridization with each examined setting. For example, the library pool that proceeded with the single-capture setting resulted in a pair of sequencing datasets named single_Mock-cycle20 and single_St2-cycle20.

To evaluate the effect of diluting the targeted gene, mock DNA was pooled with the pTAC-2 cloning vector (BioDynamics Laboratory) to vary the *amoA* gene concentration. The molar concentration ratios of mock DNA to pTAC-2 DNA were adjusted to 2:1000 (named Mock-copy1e6), 2:10000 (Mock-copy1e5), and 2:100000 (Mock-copy1e4). The final concentrations were approximately equivalent to 10^6^, 10^5^, and 10^4^ *amoA* gene copies per ng of DNA, where the lengths of the *amoA* genes, pTAKN-2, and pTAC-2 were 653–846, 2739, and 2786 bp, respectively. Sequencing libraries for the three samples were prepared as described above. For probe capture enrichment, 1 μg of library pool composed of the three equalmol libraries was used. Hybridization capture was then performed with single-capture settings to obtain capture samples but without this step for non-capture samples. Post-capture LM-PCR amplification for the Mock-copy1e6, Mock-copy1e5, and Mock-copy1e4 samples was performed for 16, 18, and 20 cycles, respectively, to obtain a sufficient amount of DNA for sequencing. The subsequent steps were performed as described above.

For the metatranscriptomic seawater samples, 20 and 19 equal-molar cDNA libraries from each transect (ON and OT, respectively) were mixed separately as a library pool, followed by hybridization capture. Evaluation using mock samples suggested that the single-capture setting was optimal for obtaining sequencing reads that reflected the original community structure (see Results). Thus, each 1 μg of the library pool proceeded using the single-capture settings as capture samples but not as non-capture samples. The sample name represents the library preparation type, sampling station, and water depth; for example, ‘capture_OT6-200m’ represents the sample collected from OT6 station at 200 mbsl layer and prepared with probe capture enrichment method, and ‘non-capture_ON1-B-5m’ represents the sample collected from ON1 station at the layer 5 m above the seafloor and without probe capturing.

For library size selection, DNA fragments <150 bp in length were removed using the GeneRead Size Selection Kit (QIAGEN) according to the manufacturer’s instructions. The final libraries of each pool were sequenced (2 × 300 bp paired-end reads) using the Illumina MiSeq sequencing platform (Illumina, San Diego, California, USA) at the Bioengineering Lab (Atsugi, Kanagawa, Japan).

### Amplicon sequencing

To compare the results with those of probe capture enrichment and shotgun sequencing methods, *amoA* amplicon sequencing was performed. The most frequently used *amoA* primer set (Arch-amoAF:5ʹ-STAATGGTCTGGCTTAGACG-3ʹ, and Arch-amoAR:5ʹ-GCGGCCATCCATCTGTATGT-3ʹ) (Francis et al., 2005), hereafter referred to as ‘Francis primers,’ was used for the amplification. A metagenomic Mock sample and three metatranscriptomic seawater samples were selected for long-read *amoA* amplicon sequencing, and the other 34 metatranscriptomic seawater samples were subjected to short-read amplicon sequencing.

The *amoA* amplicon was prepared using two-step PCR method. For the metatranscriptome samples, cDNA was prepared using the PrimeScript II 1st Strand cDNA Synthesis Kit (TaKaRa Bio). The first targeted PCR was performed using a KAPA HiFi HotStart ReadyMix PCR Kit (Roche, Basel, Switzerland). Thermal cycles were as follows: 95 °C for 2 min, (98 °C for 20 s, 65 °C for 15 s, 72 °C for 30 s) cycle for 35 times, 72 °C for 5 min. The products were then subjected to a second barcoding PCR to add index sequences, using the same PCR kit. Next, the amplicons were subjected to single-molecule real-time (SMRT) sequencing. Multiplexed amplicon libraries of *amoA* were prepared according to the ‘Amplification of bacterial full-length 16S gene with barcoded primers’ protocol (Pacific Biosciences of California, Menlo Park, California, USA). The final SMRT libraries were sequenced using a PacBio Sequel IIe system (Pacific Biosciences of California) in circular consensus sequencing (CCS) mode to generate accurate high-fidelity (HiFi) reads at the Bioengineering Lab (Sagamihara, Kanagawa, Japan).

We used 34 of 39 metatranscriptome seawater samples for short-read *amoA* amplicon sequencing, with the exception of three and two samples that were previously used or used for long-read amplicon sequencing as mentioned above. cDNA was synthesized using a Transcriptor First Strand cDNA Synthesis Kit (Roche). Amplification was performed using the Tks Gflex DNA Polymerase Low DNA (Takara Bio). Thermal cycles were as follows: 95 °C for 5 min, (94 °C for 45 s, 53 °C for 60 s, 72 °C for 60 s) cycle for 40 times, and 72 °C for 15 min. Among the 34 cDNA samples, four were excluded from further experiments because of severely low amplicon production. The final amplicon concentrations varied between 3.4 and 93.1 nM. For a realistic comparison using metatranscriptome seawater samples, the remaining amplicons, including those with an insufficient DNA concentration for Illumina library preparation, were subjected to sequencing experiments. Multiprexed libraries were prepared using the Nextera XT DNA Library Preparation Kit (Illumina) following the manufacturer’s instructions. The libraries were subjected to short-read sequencing (2 × 300 bp paired-end reads) using the Illumina MiSeq sequencing platform (Illumina) at the Atmosphere and Ocean Research Institute, The University of Tokyo (Kashiwa, Chiba, Japan).

### Bioinformatics

For capture and non-capture sequencing reads, both ends of the reads containing low-quality bases (Phred quality score < 20) and adapter sequences were trimmed using TrimGalore (https://github.com/FelixKrueger/TrimGalore) with default settings. Sequences with low complexity or shorter than 100 bp were discarded using PRINSEQ++ (Cantu et al., 2019) with default settings, and the remaining reads were defined as quality-controlled (QC) paired-end reads. QC paired-end reads were *de novo* assembled using metaSPAdes (Nurk et al., 2017) or rnaSPAdes (Bushmanova et al., 2019) with the default options for metagenome and metatranscriptome samples, respectively. Protein-coding sequences (CDSs) in the assembled contigs were predicted using Prodigal (Hyatt et al., 2010) with ‘-meta’ setting. Next, *amoA* genes were searched using DIAMOND (Buchfink et al., 2021) with ‘--max-target-seqs 1 --evalue 1e-30 --subject-cover 50 --query-cover 50 --id 60’ settings against the CuMMO gene sequence database. CDSs of <400 bp or >1000 bp were excluded from further analyses. Chimeric sequences were also removed using VSEARCH (Rognes et al., 2016) with ‘--dn 0.5’ setting. The remaining sequences were clustered into 97% sequence similarity with >50% length coverage using MMseq2 (Steinegger & Söding, 2017) with ‘--cov-mode 0’ setting. The 97% threshold has been widely used for *amoA* OTU definition (Fabien et al., 2022) and hence, we used it in this study. After discarding singletons, each cluster was designated as an operational taxonomic unit (OTU). As similar to the typical 16S rRNA amplicon sequencing analysis, we used the ‘sample concatenating’ approach: all the retrieved sequences from multiple samples were pooled for clustering to define OTU. A phylogenetic tree of the OTUs was constructed using MAFFT (Katoh & Standley, 2013) for multiple alignments and FastTree2 (Price et al., 2010) for phylogenetic tree estimation using the maximum-likelihood method with the GTR + G model, which was selected using MEGA X (Kumar et al., 2018) based on the Bayesian information criterion (BIC). The taxonomy of AOA OTUs was assigned using DIAMOND (Buchfink et al., 2021) against the AOA *amoA* sequence database defined by Alves et al. (Alves et al., 2018) after manual curation. Ecotypes of AOA OTUs were estimated based on taxonomy and phylogeny according to previous definitions (Beman et al., 2008; Francis et al., 2005). The functions of the encoding genes and the original taxonomy of each OTU were estimated via annotation using DIAMOND (Buchfink et al., 2021) against the CuMMO gene sequence database and phylogenetic tree topology. For abundance estimation, the QC paired-end reads were merged using FLASH (Magoč & Salzberg, 2011) with default settings and subject to the read mapping to OTUs using Bowtie2 (Langmead & Salzberg, 2012) with ‘-N 1’ setting for accepting single sequencing error in seed matching. The coverage of QC-merged reads was estimated using Nonpareil3 with default settings (Rodriguez-R et al., 2018).

For long-read amplicon sequencing reads, CCS reads containing at least three full-pass sub-reads on each polymerase read with >99% average base-call accuracy were retained as HiFi reads using the standard PacBio SMRT software package with the default settings. Reads of <400 bp or >1000 bp were removed for further analyses. To remove the primer sequences, 20 bp in both terminals of the HiFi reads was trimmed using SeqKit (Shen et al., 2016), and the remaining reads were defined as QC HiFi reads. Chimeric sequences were removed using VSEARCH (Rognes et al., 2016) with ‘--dn 0.5’ setting. The remaining reads were mapped to OTUs using Bowtie2 (Langmead & Salzberg, 2012) to calculate the RPKMS as described above.

For short-read amplicon sequencing reads, quality filtering was conducted using the same methods as those for capture and non-capture sequencing reads as described above. The sequencing reads (2 × 300 bp paired-end reads) were expected to not cover the entire amplicon products; the Francis primer was designed to amplify the 635 bp region of *amoA* gene (Francis et al., 2005). Therefore, for chimeric read prediction, the QC paired-end reads were concatenated with 50 bp length ‘N’ nucleotids as a linker and subjected to VSEARCH (Rognes et al., 2016) with ‘--dn 0.5’ setting. Predicted chimeric reads were removed from the QC paired-end reads using SeqKit (Shen et al., 2016). For abundance estimation, QC paired-end reads after chimera removal were directly mapped to OTUs using Bowtie2 (Langmead & Salzberg, 2012) with ‘-N 1’ setting. Only reads whose ends were aligned to the same OTU were used for abundance estimation.

The mapped reads were used to calculate the number of reads per kilobase of genes per million reads sequenced (RPKMS) for each OTU. Rarefaction curves were calculated using the vegan package with default settings after excluding samples from which <100 reads were mapped to the OTUs. The RPKMS abundance in each ecotype, along with the coordinates and water depths, were visualized using Ocean Data View (https://odv.awi.de/). Spatial interpolation was performed using a data-interpolating variational analysis (DIVA) gridding algorithm (Troupin et al., 2012).

### Data availability

All Illumina short-read and PacBio HiFi read sequences were deposited in the DDBJ Sequence Read Archive (DRA) (Table S2). All data were registered under the BioProject ID PRJDB15057 [https://ddbj.nig.ac.jp/resource/bioproject/ PRJDB15057]. The designed capture probes are available in HyperDesign (Roche Sequencing Solutions, Santa Clara, CA, USA) under the Custom Design Name 16TKYKT05.

## Results

### Validation using metagenomic mock community samples

To evaluate the efficiency of the probe capture enrichment approach, we prepared an artificial metagenomic mock DNA sample as the ground truth. The mock sample comprised six cloning vectors into which different *amoA* genes were inserted at equal molar concentrations. The *amoA* genes were selected from three AOAs and three AOBs belonging to phylogenetically distant lineages (see Materials and Methods). Using the mock sample, we sought good experimental settings for hybridization capture and post-capture LM-PCR conditions that could affect the efficiency of probe capture enrichment. OTUs at a 97% sequence identity threshold were *de novo* identified from the sequence data produced using five different experimental settings. Additionally, we performed *amoA* amplicon sequencing using the most frequently used *amoA* primer set, Francis primers (Francis et al., 2005) for comparison with the probe-capture enrichment approach.

Using our methodology (Figure 1), seven OTUs were predicted from a pool of four capture and one non-capture samples (Figure 2A, Table S3). Among the seven OTUs, all six *amoA* sequences in the cloning vectors were successfully retrieved with complete sequence matches. Only one of the other OTUs (OTU_mock6) showed high sequence similarity to those from marine uncultured AOA (Thaumarchaeota archaeon casp-thauma1), possibly indicating unexpected mixed contamination in the sample before hybridization. OTU_mock6 represented up to 0.09% of the reads in the four capture samples, suggesting a low abundance of the contaminated *amoA* gene in the samples. The mapped ratio of sequence reads to OTUs was 15.8% in the non-capture sample (non-capture_Mock). This value was as expected; the ratio of nucleotide bases belonging to *amoA* genes in the construct was close to 21%, where the lengths of *amoA* genes and the cloning vector were 651–846 and 2739 bp, respectively. In contrast, the ratios from the capture samples were higher: the ratio from a singlecapture (single_Mock-cycle20) was 72.2%, whereas those from doublecapture (double_Mock-cycle5/20, double_Mock-cycle7/14, and double_Mock-cycle7/20) ranged from 75.4 to 79.8%. In contrast to the OTU mapping analysis, almost all quality-controlled (QC) merged reads (98.4–99.7%) were mapped to the cloning vector sequences, and most reads (97.8–99.9%) showed sequence similarity to any of the six *amoA* gene sequences (blastn search with e-value <1E-5), indicating that these reads contain partial *amoA* gene sequences. These findings suggest that probe hybridization capture selectively enriched libraries containing the *amoA* gene sequences, whereas some sequencing reads failed to map to the OTUs, most likely due to the partial involvement of *amoA* gene sequence at the edge of reads that escaped OTU mapping. Almost all long-read amplicon sequencing reads were mapped to the OTUs (99.9%, amplicon_Mock).

**Figure 1.**
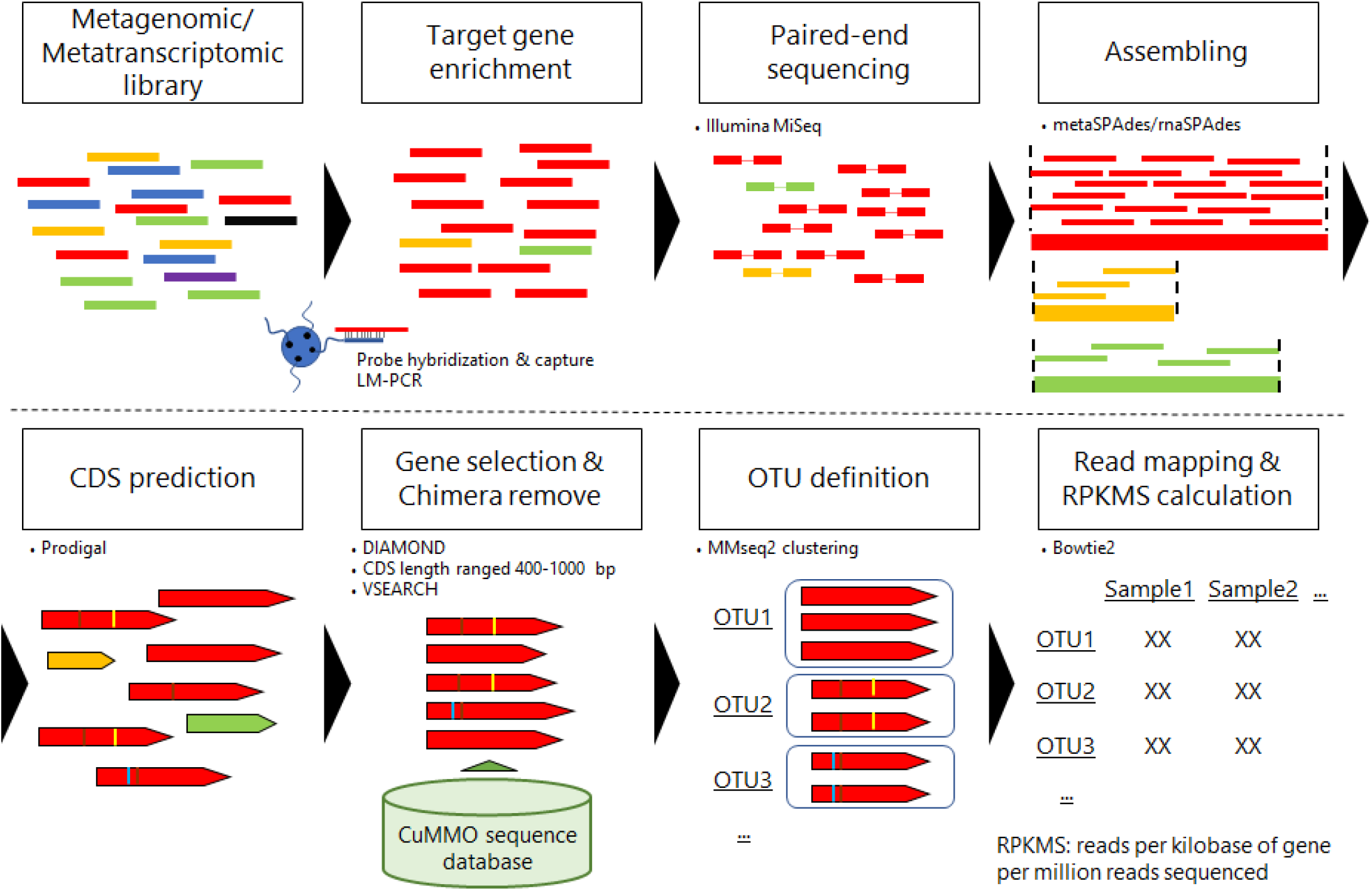
A schematic overview of probe capture enrichment sequencing analysis. The DNA/RNA extracted from the metagenomic/metatranscriptomic samples was subjected to conventional library preparation for short-read sequencing. After LM-PCR amplification (pre-capture LM-PCR), the DNA/cDNA libraries were hybridized to probes and captured using beads. The recruited libraries are LM-PCR amplified (post-capture LM-PCR) and sequenced using the Illumina MiSeq platform. Sequencing reads after quality control (QC) are assembled using metaSPAdes or rnaSPAdes for metagenomic and metatranscriptomic samples, respectively. Protein coding sequences (CDSs) are predicted using Prodigal software. The CDSs that showed sequence similarity to CuMMO genes using DIAMOND, with expected nucleotide lengths (400–1000 bp), and without chimeric sequences estimated using VSEARCH, were clustered with >97% thresholds using MMseq2 for the operational taxonomic unit (OTU) definition. QC reads were mapped to the OTU sequences using Bowtie2 to calculate the reads per kilobase of genes per million reads sequenced (RPKMS) to estimate the abundance of each OTU in the samples.

**Figure 2.**
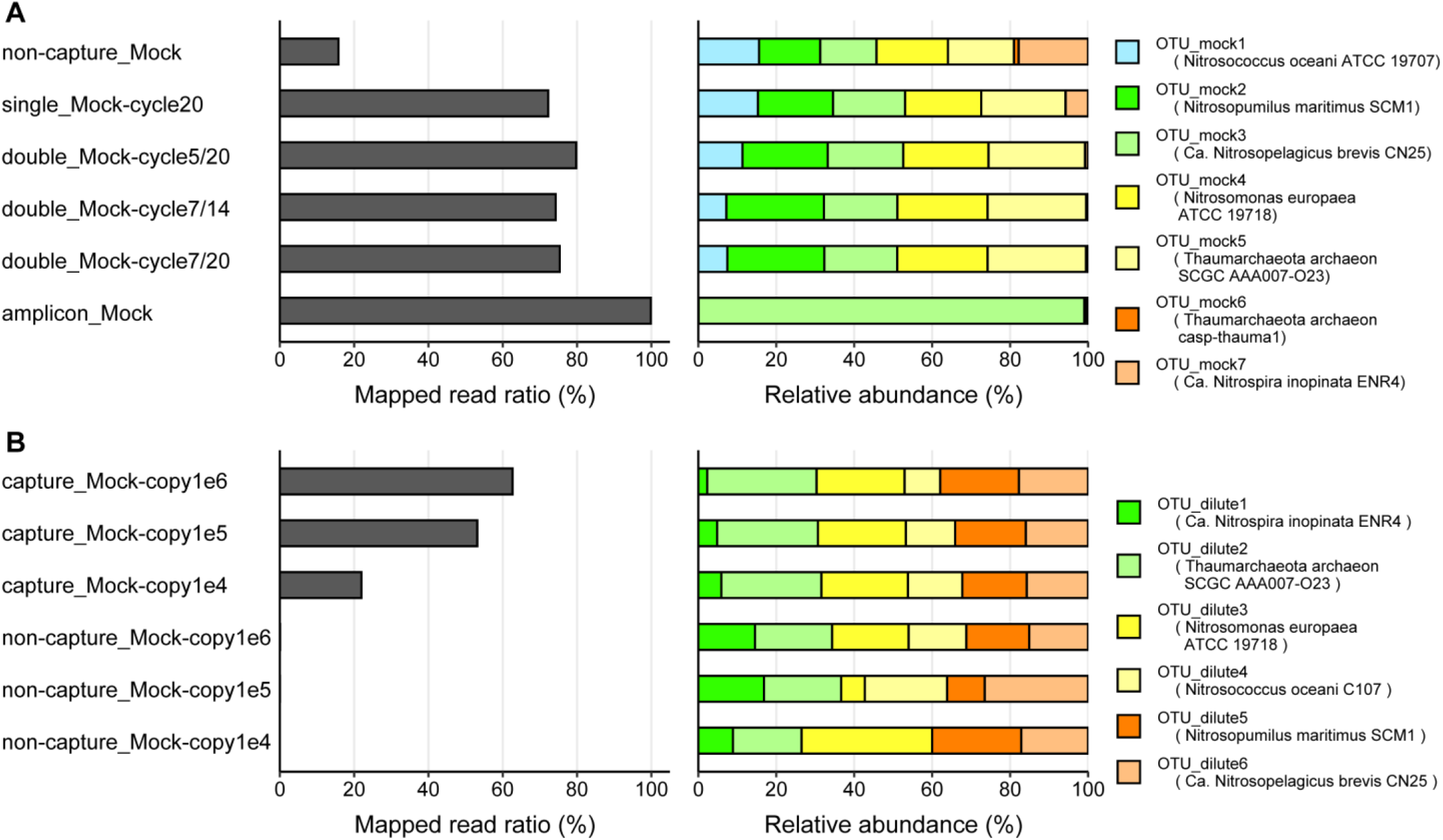
The *amoA* probe capture enrichment sequencing of metagenomic mock samples. The mock sample was comprised of six cloning vectors containing artificially synthesized *amoA* genes from three AOA and three AOB lineages at the same molar concentration. The left bar represents the ratio of mapped reads to OTU sequences. The right percentage-stacked bar plot represents the relative abundance of each OTU based on RPKMS. Lineages assigned to each OTU are shown in the figure legends. (**A**) Probe capture enrichment was performed using different numbers of hybridization capture repetitions and post-capture LM-PCR cycle settings. Amplicon sequencing was performed using the Francis primers. OTU were defined using pooled reads from all five capture samples; that is, those from amplicon sequencing were not used. (**B**) Diluted samples were prepared using the original mock sample rarefied with control DNAs (cloning vector pTAC-2) and adjusted to 10^6^, 10^5^, and 10^4^ *amoA* gene copies per ng of DNA. Data generated with and without probe capture enrichment are represented as ‘capture’ and ‘non-capture’, respectively, in the sample name. OTU were defined using the pooled reads from all six samples.

The relative abundance of reads per kilobase of genes per million reads sequenced (RPKMS) of the six OTUs corresponding to the inserted *amoA* genes was generally consistent under non-capture settings, but fluctuated slightly in the capture samples. This trend was attributed to a decrease in the OTU_mock7 (*Ca*. Nitrospira inopinata ENR4), with abundances of 5.7% and 0.6% in the single- and double-capture settings, respectively. Similarly, OTU_mock1 (*Nitrosococcus oceani* ATCC 19707) showed a slight decreasing tendency with an increase in the probe hybridization capture step. This may be explained by the lower recruitment power of the probes against the OTUs; among the CuMMO gene sequences used for probe design, only 3 and 39 sequences showed similarity to the *amoA* genes from *Ca.* Nitrospira inopinata ENR4 and *Nitrosococcus oceani* ATCC 19707, respectively, and more than 100 sequences from the other 4 organisms (blastn search with identity >95% and e-value <1E-30 criteria). Considering that probe hybridization accepts a few mismatches, the uneven sequence distribution in probe design may have caused a bias in probe capture enrichment. Only one OTU (OTU_mock3; *Ca.* Nitrosopelagicus brevis CN25, which belongs to the WCA ecotype (Santoro et al., 2015)) occupied almost all parts of the amplicon sequencing sample (99.0%, amplicon_Mock). This was likely due to a primer bias; the primer set used for amplification (Francis primers) perfectly matched the *amoA* sequences from *Ca.* Nitrosopelagicus brevis CN25 possessed three and two mismatches with the other two AOA *amoA* sequences (*Nitrosopumilus maritimus* SCM1 and Thaumarchaeota archaeon SCGC AAA007-O23, respectively), and showed no significant match with any of the three AOB *amoA* sequences. Collectively, despite the potential slight bias in some ammonia oxidizers, the probe capture enrichment strategy effectively enriched *amoA* genes with low compositional changes and high sensitivity, overwhelming metagenomic and amplicon sequencing approaches.

### Study of targeted gene concentration sensitivity

Next, we evaluated the effect of the target gene concentration on the probe capture enrichment efficiency. Diluted metagenomic mock samples were prepared, in which the total concentrations of the six *amoA* genes were adjusted to 10^6^, 10^5^, and 10^4^ copies/ng of DNA using commercially available plasmid DNA, referred to as Mock-copy1e6, Mock-copy1e5, and Mock-copy1e4, respectively. The adjusted concentrations were compatible with those in a wide range of environments such as soil (Bahram et al., 2022; Bannert et al., 2011; Ding et al., 2020; Han et al., 2013; Marusenko et al., 2013), wastewater (Spasov et al., 2020; Wang et al., 2021; Zhang et al., 2015), estuaries (Hollibaugh et al., 2011), and seawater (Christman et al., 2011). OTUs were *de novo* predicted using pooled reads from three capture and three non-capture samples. Using probe capture enrichment and our bioinformatic methodology with ‘single-capture’ settings against diluted samples, all six induced *amoA* genes were successfully identified (Figure 2B). The mapped read ratios on the OTUs decreased with the dilution rate; 63.1%, 53.7%, and 22.3% were mapped from samples with 10^6^, 10^5^, and 10^4^ copies per ng DNA (capture_Mock-copy1e6, capture_Mock-copy1e5, and capture_Mock-copy1e4, respectively). In contrast, only 0.0–0.1% of the reads were mapped from the non-capture samples (non-capture_Mock-copy1e6, non-capture_Mock-copy1e5, and non-capture_Mock-copy1e4). When comparing the ratios between samples with the same *amoA* concentration, the capture samples showed 990–8100 times higher values than the non-capture samples. The relative RPKMS abundances of the OTUs were generally even within the capture samples, whereas they were disturbed in the non-capture samples because of the extremely low numbers of mapped reads. Overall, the results demonstrate the high efficiency and specificity of our probe capture enrichment approach under conditions of low target gene concentrations, analogous to the natural environment.

### Study of enrichment settings for metatranscriptomic seawater samples

To elucidate the efficiency of the probe capture enrichment approach for metatranscriptomic data and the potential impact of post-capture LM-PCR settings on performance, we analyzed cDNA libraries prepared using a seawater sample collected at station St2. Using the pooled reads from four samples with different settings, eight OTUs were identified (Figure S2). The mapped read ratios were low in the single-capture sample (0.5%, single_St2-cycle20) and original metatranscriptomic sample (0.1%, non-capture_St2). The numbers of sequencing reads from non-capture and single-capture samples were 2.3 M and 2.2 M, respectively. The relative abundances were concordant between the two samples, suggesting a low compositional effect in single-capture enrichment settings. Notably, a comammox member (OTU_st2-8; assigned to *Ca.* Nitrospira inopinata), which was hardly amplified using the Francis primers (Daims et al., 2015; van Kessel et al., 2015). In contrast to the non-capture and single-capture samples, the double-capture samples showed higher mapped read ratios (86.0% and 92.6% for double_St2-cycle5/20 and double_St2-cycle7/14, respectively) (Table S3). In addition, likely because of the severely low number of sequencing reads, RPKMS relative abundances in double_St2-cycle7/14 were disturbed compared with those in the other samples. Taken together, the analysis demonstrates that single-capture settings are applicable to marine metatranscriptomic samples, whereas double-capture settings can produce unstable results.

### Probe capture enrichment sequencing of metatranscriptomic seawater samples

Using the single-capture settings described above, we conducted probe capture enrichment sequencing of *amoA* genes against 39 metatranscriptomic seawater samples collected at 11 stations spanning from the coast to the open ocean at different water depths (Figure S1, Table S2). In addition to the capture and non-capture samples, we performed short- and long-read *amoA* amplicon sequencing of 34 and 3 metatranscriptomic seawater samples, respectively, using Francis primers for comparison.

Despite the equalmol libraries in a pool before the probe hybridization capture, the numbers of sequencing read from each captured sample ranged from 0.29 to 3.28 M (1.25 ± 0.82 M on average) with an increasing tendency along with water depth and distance from land (Table S3). In addition, among the seawater samples used for *amoA* amplification, no obvious amplicons were obtained via PCR amplification from four samples (ON3-0m, ON4-0m, St2-0m, and OT4-0m), all of which were collected from the most surface layer at each station. These results reflect the general tendency of ammonia oxidizers to be more abundant in the deeper layers than in the surface layers and outside the bay rather than inside the bay, as widely observed previously (Lu et al., 2020; Molina et al., 2020; Newell et al., 2013; Shafiee et al., 2021; Tolar et al., 2013; Zou et al., 2020). This tendency was also concordant with the geochemical analysis, which showed that nitrate (NO_3_ ^-^) and ammonium (NH_4_ ^+^) concentrations were lower in the samples collected at the surface than near the bottom, and in the coastal areas than in the pelagic areas (Table S2). In contrast, as expected, the read numbers were generally even (0.96–1.89 M, 1.19 ± 0.17 M on average) within the non-capture samples. Notably, among the three long-read amplicon sequencing samples, 34.2% (amplicon_OT3-50m) and 34.2% (amplicon_OT3-B-5m) of the data were lost, mainly due to chimeric read filtering (Table S3), indicating drastic chimeric production during the amplification step. A similar tendency was also observed in the short-read amplicon sequencing, where 35.6 ± 5.4% of reads were removed in this step (Table S3). The estimated coverage of the sequencing reads in all capture and non-capture samples was high (>91.7%), suggesting sufficient sequencing depth to cover the metatranscriptomes overall (Figure S3A).

The probe capture enrichment approach revealed the expression levels of *amoA* genes by ammonia oxidizers in the community at high resolution (Figure 3). We identified 87 OTUs, of which at least 78 and 2 OTUs were assigned to *amoA* genes from AOA and AOB, respectively. Among the other OTUs, two and four were estimated to be particulate methane monooxygenase (*pmoA*) and Cytochrome P450 genes, respectively, from methane-oxidizing bacteria (MOB) and Actinomycetota, both of which are included in CuMMOs and are concordant with the designed probe (see Materials and Methods). The other OTU showed low (84.8%) sequence similarity to known *amoA* genes, as described by Alves et al. (Alves et al., 2018), and considering the phylogenetic topology, it was difficult to conclude whether AOA *amoA* or functionally different gene. In contrast, possible 78 AOA OTUs showed 94.2–100% sequence similarity to *amoA* genes in the sequence database (Alves et al., 2018). Mapped read ratios were significantly higher for the capture samples (0.8%–82.0%, 50.4 ± 27.2% on average) than for non-capture samples (0.02%–0.08%, 0.05 ± 0.02%) (p < 0.05, Mann–Whitney U-test [U-test]). Almost all the reads (>99.3%) from the long-read amplicon sequencing corresponded to the OTUs, whereas the mapped read ratio from the short-read amplicon sequencing was moderate and fractuated (0.13%–40.0%, 20.6 ± 13.6% on average) likely because of the challenging amplification and chimera detection for paired-end reads (see Materials and Methods). In addition, the number of detected OTUs in the capture samples (ranging from 35 to 85, 61.3 on average) was significantly higher than that in the non-capture samples (ranging from 4 to 27, 11.3 on average) (p < 0.05, U-test, Bonferroni correction). Those from amplicon sequencing (ranging from 2 to 25, 8.2 on average) were significantly higher than those from the non-capture samples, whereas they were significantly lower than those from the capture samples (p < 0.05, U-test, Bonferroni correction), and no AOBs were detected. Among the AOA and AOB OTUs, 36 (41%), 68 (77%), and 54 (61%) were unidentified in the non-capture, long-read amplicon, and short-read amplicon sequencing samples, respectively, whereas 30 (34%) OTUs were not detected in any of the non-capture and amplicon sequencing samples. In contrast to the capture samples, the rarefaction curves of most non-capture and amplicon sequencing samples were not saturated, with very low numbers of reads mapped to OTUs, indicating that more sequencing reads are required to cover the AOA and AOB populations in the communities (Figure S3B).

**Figure 3.**
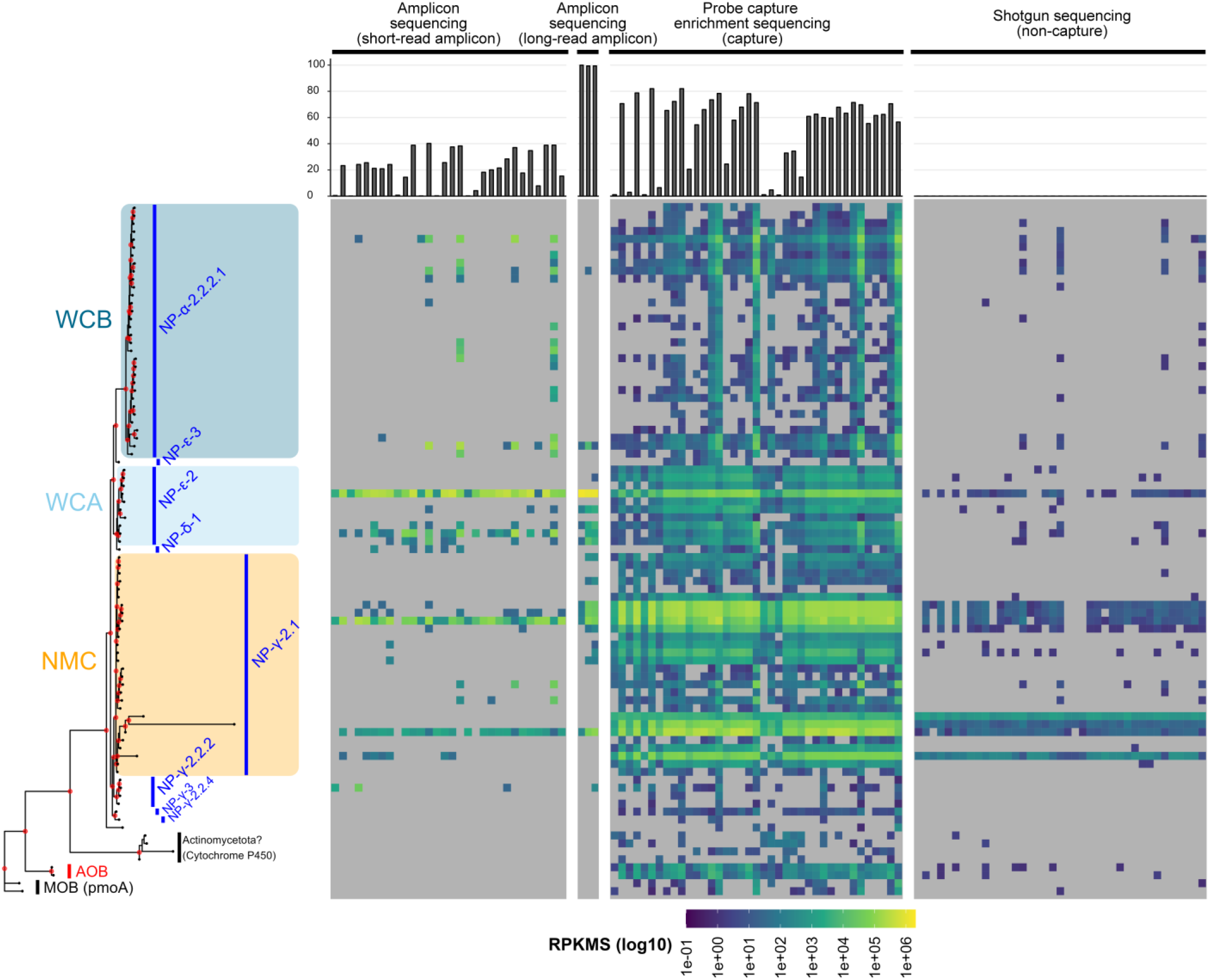
The *amoA* amplicon, probe capture enrichment, and shotgun sequencings of metatranscriptomic seawater samples. OTUs were predicted using pooled reads from all 39 capture and 39 non-capture samples, excluding those from 30 and 3 short- and long-read amplicon sequencing samples, respectively. For amplicon sequencing, *amoA* was amplified from the metatranscriptomic cDNA using Francis primers. The left panel represents a phylogenetic tree of the OTUs obtained using the maximum-likelihood method. External nodes were grouped into clades defined by Alves et al. for ammonia-oxidizing archaea (AOA) (blue bars and text), the domain level for ammonia-oxidizing bacteria (AOB) (red), methane-oxidizing bacteria (MOB), and possible Actinomycetota with genes other than *amoA* (black). The background color indicates the three major AOA ecotypes in the ocean. The red dots at the internal nodes indicate support values of >80%. The top bar plot represents the ratio of mapped reads to OTUs by the total QC merged or QC HiFi reads. The heat maps represent the RPKMSfor each OTU in each seawater sample. The heat maps were separated into short-read amplicons, long-read amplicons, and capture, and non-capture samples. Gray cells indicate OTUs to which no reads were mapped.

All the AOA OTUs were assigned to order Nitrosopumilales and spanned to four order-level clades (NP-α, NP-δ, NP-ε, NP-γ). NP-α and NP-ε were reported to be observed in pelagic marine water, whereas NP-δ was both coastal water and marine sediment, and NP-γ was composed of members inhabiting all three oceanic environments (Alves et al., 2018), as consistent with the samples (Figure S1, Table S2). The NMDS analysis showed a significant association of the composition with water depth (p < 0.05, permutation tests for regression models implemented by ‘envfit’ function, vegan package) (Figure S4). Especially, in the capture samples, expression levels of WCB (OTUs assigned to NP-α-2.2.2.1) were increased along with water depth, whereas NMC (NP-γ-2.1) were decreased and WCA (NP-ε-2) were evenly distributed (Figure 4). Overall, the NMC was the dominant group, followed by the WCB and WCA. These findings were scarcely observed in non-capture samples. In contrast, to capture and non-capture sequencing, the WCA was the most dominant ecotype in the expression profiles from amplicon sequencing, which was likely due to PCR bias caused by the Francis primers, as described above. These results are consistent with those of previous studies that investigated the AOA community structures in the ocean using *amoA* amplicon sequencing (Ijichi et al., 2019; Lu et al., 2019; Santoro et al., 2017; Shiozaki et al., 2016; Wuchter et al., 2006).

**Figure 4.**
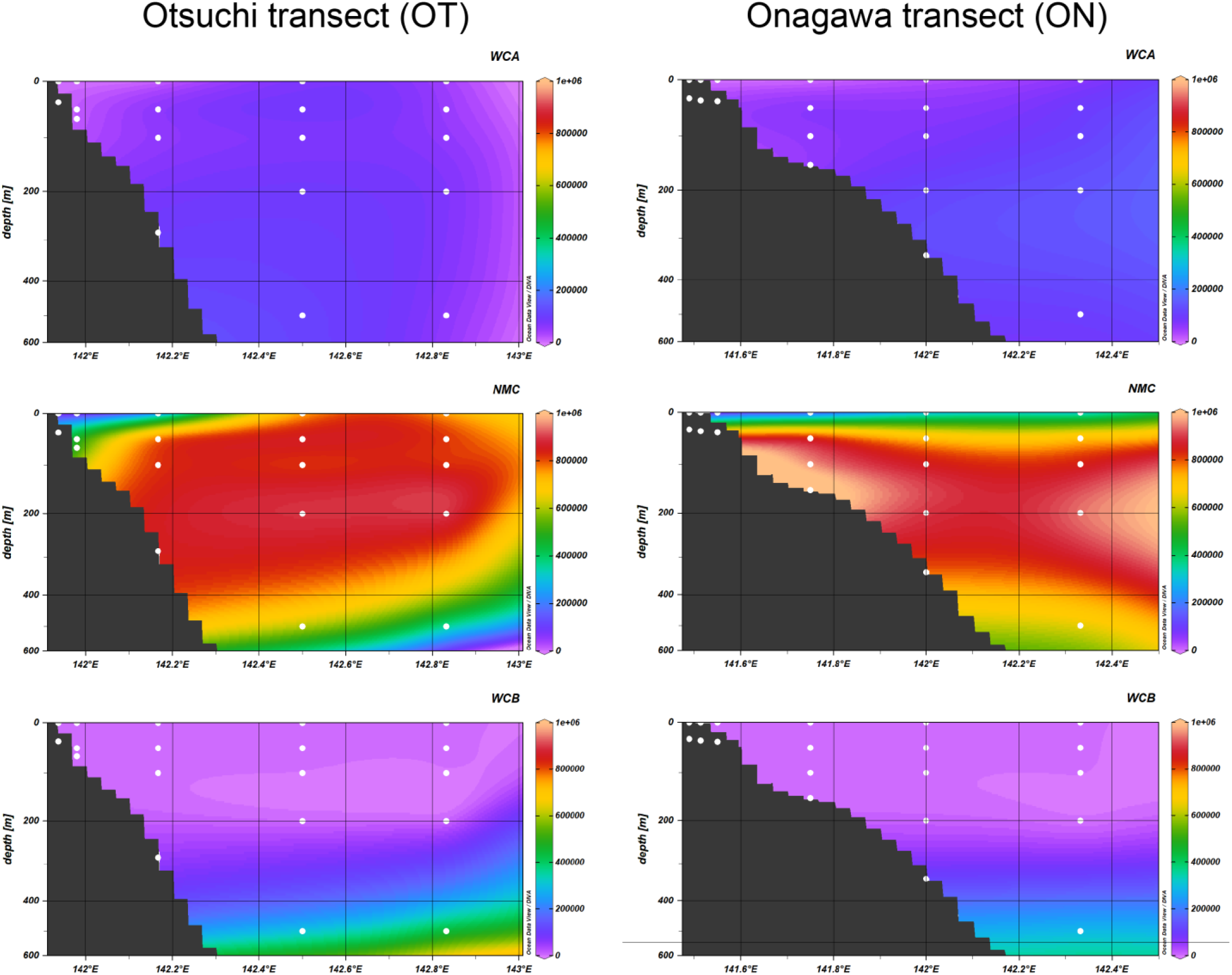
RPKMS abundances of the three major ecotypes in the ocean along the east-west section of Onagawa (ON) and Otsuchi (OT) transects. White dots represent the sampling depths at each station. WCA, NMC, and WCB are represented by OTUs assigned to NP-ε-2, NP-γ-2.1, and NP-α-2.2.2.1 (see Figure 3), and total RPKMS values in each ecotype were used in this analysis.

## Discussions

We developed a probe-capture enrichment sequencing method for the *amoA* gene to effectively investigate the community structure of ammonia oxidizers and their potential activities (Figure 1). By applying this approach to metagenomic mock community samples, our method successfully reconstructed the true community structure, demonstrating that the approach can provide a qualitatively reliable estimation of AOAs and AOBs composition in community samples. In addition, this approach successfully identified *amoA* expression levels of diverse ammonia oxidizers in oceanic samples with high phylogenetic resolution. This performance was contrary to that of shotgun metatranscriptome and amplicon sequencing, both of which resulted in highly divergent profiles with lower resolution. Chimeric sequence removal or sequence clustering for OTU definition did not cause a severe reduction in mapping efficiency (Figure 1, Table S4), suggesting that the principal *amoA* gene sequences were retrieved as OTUs. On the contrary, chimeric read filtration from amplicon sequencing deleted more than 30% of the reads on average (Table S3). This finding emphasizes the efficacy of the PCR-free approach. Taken together, probe capture enrichment sequencing provides detailed and comprehensive information on *amoA* genes in the prokaryotic community, thereby contributing to a substantial understanding of the prokaryotic nitrogen cycle in the environment.

In contrast to 16S rRNA genes, which contain nine hypervariable regions (V1-V9) flanked by highly conserved sequences throughout the bacterial and archaeal domains, there are no well-conserved regions in most functional genes, including *amoA*. This severely restricts the design of a ‘universal’ primer set that perfectly matches diverse *amoA* genes covering AOAs and AOBs. Thus, common *amoA* amplicon sequencing can underestimate many lineages of ammonia oxidizers because of primer mismatches, and in contrast, it will overestimate specific members who possess genes with higher binding affinity to the primer. Our experiments revealed a significant PCR bias in the *amoA* gene from *Ca.* Nitrosopelagicus brevis CN25, a member of the WCA family(Figure 2). The same trend was observed in the field metatranscriptomic samples: WCA was predominantly detected by amplicon sequencing, whereas NMC was more abundant by probe capture enrichment and simple shotgun sequencing (Figure 3). Current knowledge of ecotypic distribution in the ocean is thus likely to be affected by primer biases, and further reconsideration of the potential contribution of NMC as well as WCA to the nitrogen cycle in the ocean is needed. Determining the spatial distribution and temporal variation of the clades, particularly in coastal zones with oceanographically dynamic areas (Shiozaki et al., 2015), necessitates further large-scale sampling and analysis.

Meta-omics (i.e., metagenomics and metatranscriptomics) relies on shotgun sequencing and is now widely used to investigate the phylogenetic and genomic diversity of AOAs and AOBs, as well as the potential metabolic pathways involved in the nitrogen cycle. One advantage of this approach is that it avoids PCR bias, allowing for the recovery of sequences that would escape conventional PCR amplification (Cheung et al., 2019; Delmont et al., 2018). In the present study, OTUs that were not detected by amplicon sequencing were identified by probe capture enrichment and simple shotgun sequencing (Figure 3). Using a shotgun sequencing approach, it was recently reported that ammonia oxidizers in the ocean are more diverse than previously thought (Cheung et al., 2019). However, as we have demonstrated, the low concentration of target genes/transcripts in the heterogeneous community severely limits the efficiency of this approach. Here, the probe capture enrichment approach showed a greater number and more diverse expression levels of *amoA* than simple shotgun sequencing. This trend is similar to that observed in a previous study focusing on 16S rRNA (Gasc & Peyret, 2018). Consequently, the probe capture enrichment approach provides more informative results for *amoA* sequences than the typical omics-based methods.

Although a few recent studies have made great efforts toward the taxonomic classification of diverse *amoA* genes(Alves et al., 2018), there is still no comprehensive *amoA* gene sequence database including those from both AOAs and AOBs. This is in sharp contrast to the 16S rRNA genes for which databases such as SILVA (Quast et al., 2013) or RDP (Cole et al., 2014) have been developed and are commonly used for taxonomic classification of any 16S rRNA gene sequences of interest. A previous probe capture enrichment sequencing analysis focusing on 16S rRNA in soil microbial communities showed alignment-based direct taxonomic classification of the reads via the MG-RAST pipeline (Manoharan et al., 2015; Meyer et al., 2008), which is challenging in the case of the *amoA* gene. Therefore, we constructed de novo OTUs and used a mapping-based approach for taxonomic classification and abundance estimation. As demonstrated, this approach successfully provided good information for phylogenetic estimation and was effectively utilized for RPKMF calculations with high sensitivity. However, it should be noted that reads that straddle the edge of the targeted gene sequence (i.e., reads across the *amoA* gene and adjacent genomic region) are frequently unmapped to the OTUs owing to low alignment scores that do not exceed the defined criteria, indicating the existence of false-negative reads. Indeed, in the case of the four capture samples using metagenomic Mock samples (Figure 2), the mapping ratios of the QC merged reads to OTUs were 26–30% lower than those mapped to the plasmid constructions. In addition, most reads were predicted to possess partial *amoA* sequences based on similarity search analysis. These results suggest that probe capture successfully enriched reads containing targeted genes, whereas the mapping ratio led to an underestimation of the true recurrence power of the probe capture.

It should also be noted that this method must consider the potential bias of weak recruitment power. Multiple hybridization captures (i.e., double-capture settings) generated disturbed results, whereas single-capture settings generated low numbers of sequencing reads in some cases (Figure 2A, Table S3). These issues warrant further improvements in the method to obtain more accurate results through economic sequencing. In addition, we found a possible underestimation of some ammonia oxidizers, likely because of the low number of sequences that showed similarity to the *amoA* genes in the sequence database used for probe design, resulting in a shortage of probes that hybridized to the DNA fragments containing the *amoA* genes. (Figure 2A). To overcome this bias, it is necessary to expand the targeted gene sequences for probe design to include more diverse *amoA* genes found in nature, as well as decimate high-density sequences to avoid overdetection of specific clades, as implemented in the MetCap pipeline (Kushwaha et al., 2015). Notably, similar to numerous studies in the field of environmental microbiology, sequencing experiments were conducted once per sample in this study. Further examinations with sufficient technical and biological replicates are required to obtain statistically robust conclusions. The combination of probe capture enrichment sequencing with other techniques such as 16S rRNA gene amplicon sequencing and qPCR analysis reinforces the findings of study. The *amoA* probe capture enrichment sequencing methodology is not limited to the oceanic environment but is applicable to any community sample, including soil, freshwater, sediment, and wastewater treatment plants. Further application of this method will lead to innovative findings regarding the nitrogen cycle and microbial ecology.

## Conclusion

This study demonstrated that the probe capture enrichment sequencing approach can effectively recover the community structure of ammonia oxidizers and the expression levels of *amoA* genes in the community, outperforming conventional methods. From the application of this method to oceanic metatranscriptome samples, we detected a higher diversity of AOAs than ever before, even if their expression levels were likely low in total microbial gene expression. Importantly, from the analysis of oceanic samples, we found that amplicon sequencing incorporating the most common primer set caused a significant PCR bias (i.e., most ammonia oxidizers were missed, and selective WCA members were amplified), resulting in low phylogenetic resolution and inconsistent relative abundance profiles. This indicates that *amoA* amplicon sequencing generally produces results far from the true composition, although almost all current knowledge of the phylogenetic diversity and spatial distribution of ammonia oxidizers has been based on conventional techniques. Thus, this approach could be an alternative tool to examine the community structure and biological responses of ammonia oxidizers at high resolution to understand the geochemical contributions of nitrogen-cycling prokaryotes in vast environments on Earth.

## Acknowledgements

We would like to thank the captain, crew, and onboard scientists and technicians of the R/V Shinsei-maru (JAMSTEC) during the KS16-01 cruise. Bioinformatic analysis was performed on the supercomputing system at National Institute of Genetics (NIG), Research Organization of Information and Systems (ROIS), Japan.

## Contributions

SH designed the study, performed the bioinformatics analyses, and wrote the manuscript. MI conceived and designed the study, performed the sampling and molecular experiments, and wrote the manuscript. HT designed and performs the molecular experiments. YK contributed to the probe design. CCY contributed to the bioinformatics analyses. YMK contributed to the molecular experiments. HF performed nutrient measurements. SY, WI, and KK supervised the project. TS designed the study, performed the short-read amplicon sequencing, wrote the manuscript, and supervised the project. All authors read and approved the final manuscript.

## Ethics declarations

Minoru Ijichi is an employee of QIAGEN (Chuo-ku, Tokyo, JAPAN) and a former employee of the Bioengineering Lab (Sagamihara, Kanagawa, Japan). The remaining authors declare no conflicts of interest.

