## Supplementary Information for "Probe capture enrichment sequencing of *amoA* genes discloses diverse ammonia-oxidizing archaeal and bacterial populations"

This PDF file includes:

Supplementary Tables S1 to S5.

Supplementary Figures S1 to S4.

**Supplementary Tables**

Table S1. Summary of leading studies on the distribution of ammonia oxidizers in the ocean.

Table S2. Descriptions of sampling sites and geochemistry.

Table S3. Descriptions and statistics of sequenced reads.

Table S4. Mapping ratio of sequencing reads to sequence data at each processing stage.

Table S5. RPKMS abundance in metatranscriptomic seawater samples.

### Supplementary Figures

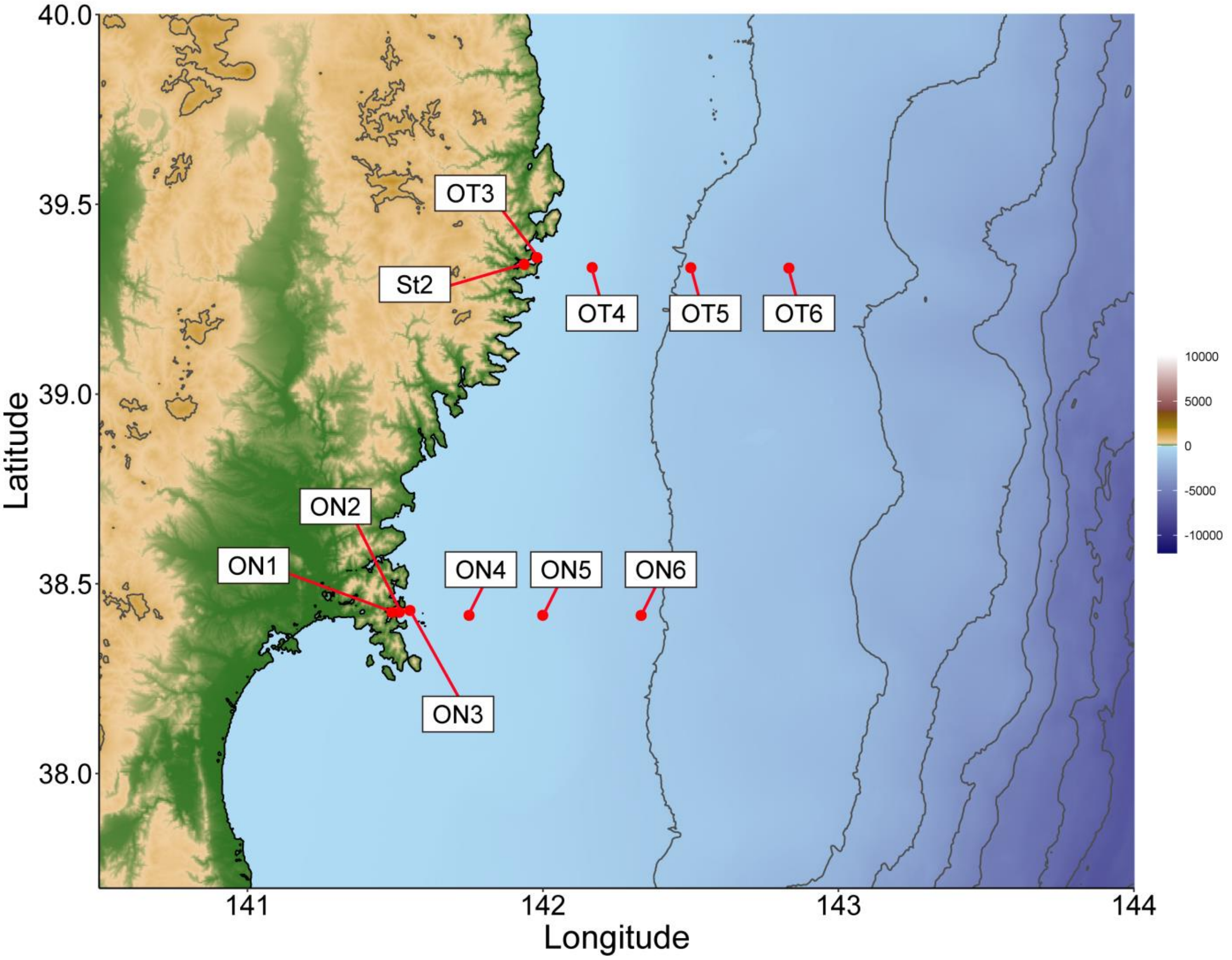


Figure S1. Map of sampling stations along the two transect lines.

The north and south lines are referred to as the Otsuchi (OT) and Onagawa (ON) transects, respectively.

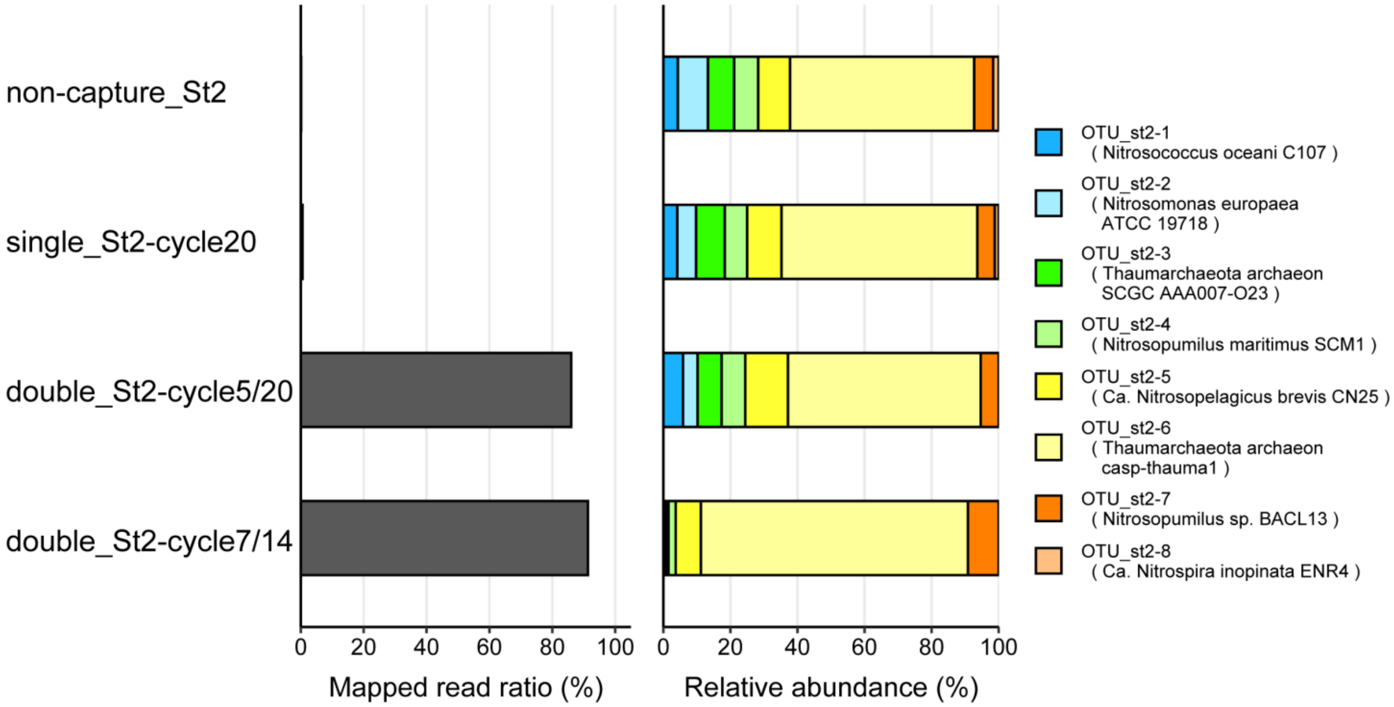


Figure S2. Analysis of metatranscriptomic seawater samples with different hybridization capture and post-capture LM-PCR settings.

The sample was collected at the 0 mbsl layer from the coastal St2 station (St2-0m). See Figure 2.

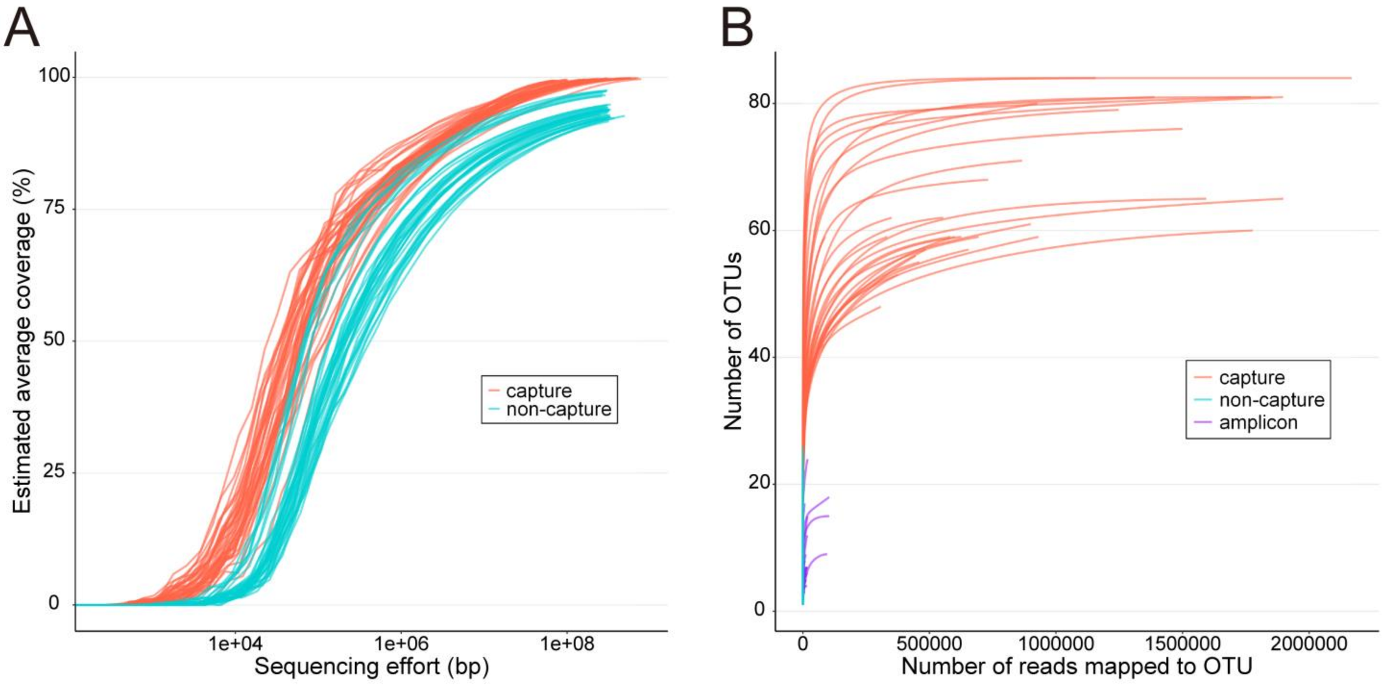


Figure S3. Estimated coverage and rarefaction curve of metatranscriptomic marine samples.

(A) Nonpareil curves of the metatranscriptomic data. Capture and non-capture samples are colored red and blue, respectively. (B) Rarefaction curve of OTU numbers against each of the examined sequencing data. Data from capture, non-capture, and amplicon sequencing settings are colored red, blue, and purple, respectively. Data with less than 100 reads mapped to the OTUs were excluded in this analysis.

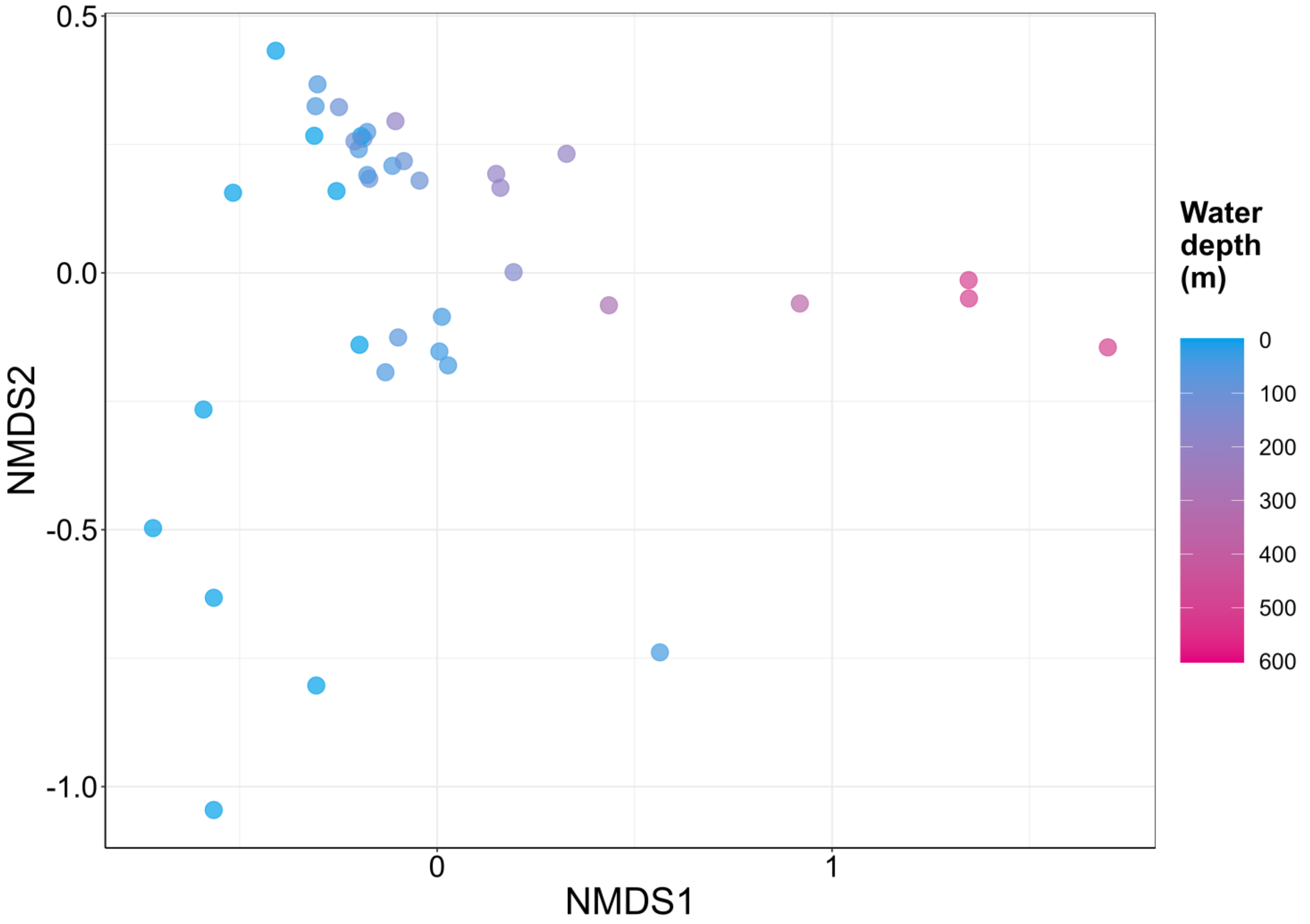


Figure S4. NMDS analysis of RPKMS abundance from metatranscriptomic seawater capture samples.
